# The Significance of Uneven Walking Transitory Modulations in Walking Momentum Regulation and Active Work

**DOI:** 10.1101/2024.07.16.603711

**Authors:** Seyed Saleh Hosseini Yazdi

## Abstract

Uneven walking presents challenges such as balance maintenance and increased energetics. Beyond average step parameter variations, humans may employ transient strategies to conserve mechanical work or enhance momentum with minimal metabolic costs. Thus, we quantified mid-flight positive mechanical work induction, step length, and effective leg length modulation from four steps before to four steps after a specific encounter for young and older adults with normal and restricted lookaheads. Simulations were also conducted to assess the impact of step length or effective leg length changes on step-to-step transition or post-transition speed. We observed that young adults’ mid-flight energy inducements were focused around the encounter, influenced by lookahead state affecting modulation based on feedback or feedforward control. While with the restricted lookahead, older adults showed similar regulation, their anticipatory modulation was poorer and extended over the evaluation interval. With normal lookahead, young adults reduced step length just before the encounter, potentially increasing momentum with less heel-strike energy dissipation. Step length changes around and after the perturbation may have been passive. Older adults exhibited longer modulation periods than young adults. We also noted active leg length modulation, likely influenced by tactile sensory information. For up-steps, effective leg lengths were shorter, while for down-steps, they were longer, potentially compensatory actions to minimize COM vertical fluctuations and associated work against gravity. The amplitude of leg length modulation may have been constrained by the flexed legs walking energetic cost or and ankle range of motion. With restricted lookahead, older adults showed larger leg length modulation amplitudes.

## Introduction

Humans traverse many walking terrains, often vastly different from even walking. Adapting gait parameters to these terrains is complex, with uneven terrains inducing perturbations that force humans to adjust their gait parameters accordingly. Some gait parameters may change on average during uneven walking, such as the reported tendency for humans to walk with shorter steps [1]. However, the step-to-step modulation may differ significantly from these average magnitudes, as they are specifically designed to regulate walking and sustain gait at each unique step. This complicated process is further explored through the investigation of Center of Mass (COM) velocity (experimental) and active works (push-off by simulation), with or without visual information about the impending perturbations. Yet, other gait parameters likely undergo modulation during the step-to-step transitions, such as effective leg length [2] or step length [3], which are currently only investigated for the point of encounter. The regulation of mid-flight energy induction, particularly when pre-emptive push-off energy exertions are inadequate, has received insufficient attention.

Prior research suggests that the lookahead’s state significantly influences humans’ strategies to navigate complex terrains. With the normal lookahead, it is proposed that humans adopt an anticipatory (feedforward) control to prepare for the terrain complexity encounters [4]. It is reported that humans accelerate [5] by exerting larger active work (push-off) [4] to attain the elevated momentum needed to compensate for the work against gravity [6]. However, the extent of the lookahead horizon to achieve a nominal walking is debated. It is indicated that humans tend to use the Just-In-Time strategy to modulate their gait [7], targeting the next proper foothold [8]. It is reported that humans tend to concentrate their gaze two to three steps ahead when walking over complex natural terrains [9]. Nevertheless, another research suggests that humans must have a six to eight-step view horizon to reach the optimal gait [10]. It is also suggested that after the point of encounter, people switch from feedforward control to feedback [4]. Thus, the vision contribution after the encounter is most likely marginal.

Contrary to the normal lookahead, with the restricted lookahead, humans cannot perform any regulatory action before the point of encounter. Nevertheless, humans do not receive all the sensory information only through vision. Vestibular [11], flight time [2], and touch [12] also provide sensory information about the walking terrain that most likely affects the step-to-step modulations regardless of the lookahead state. Any walking modulation is exerted till the associated costs outperform its benefits.

In this study, we aim to investigate the active step-to-step modulation of mid-flight work, effective leg length, and step length during uneven walking. We also seek to evaluate the influence of age and the state of lookahead on these modulations. Notably, the existing walking models [13], [14] do not consider the effects of leg length, step length, or mid-flight active work. Therefore, we cannot make predictions and compare them with experimental results. However, our primary focus is to interpret the implications of these active modulations.

## Material and Methods

We made modifications to an instrumented treadmill structure to accommodate uneven terrains (Bertec Inc. Columbus, OH, USA). These terrains were created by attaching construction foams across the width of conventional treadmill belts. The foams’ length matched the treadmill belt width, and their width was 0.0245m to allow for rotation around the treadmill rollers [1]. The foam heights were 0.019m, 0.032m, and 0.045m (Figure *1*A). We used the maximum foam heights (peak to peak) to differentiate the terrains (terrain amplitude). Each profile length was set to 0.3m, allowing for a level foot landing or an incline covering two adjacent profiles. The uneven terrains were designed to be wrapped around the original belt and closed using alligator lacing clips. Subsequently, the terrain and the original belts were re-tensioned together.

**Figure 1:**
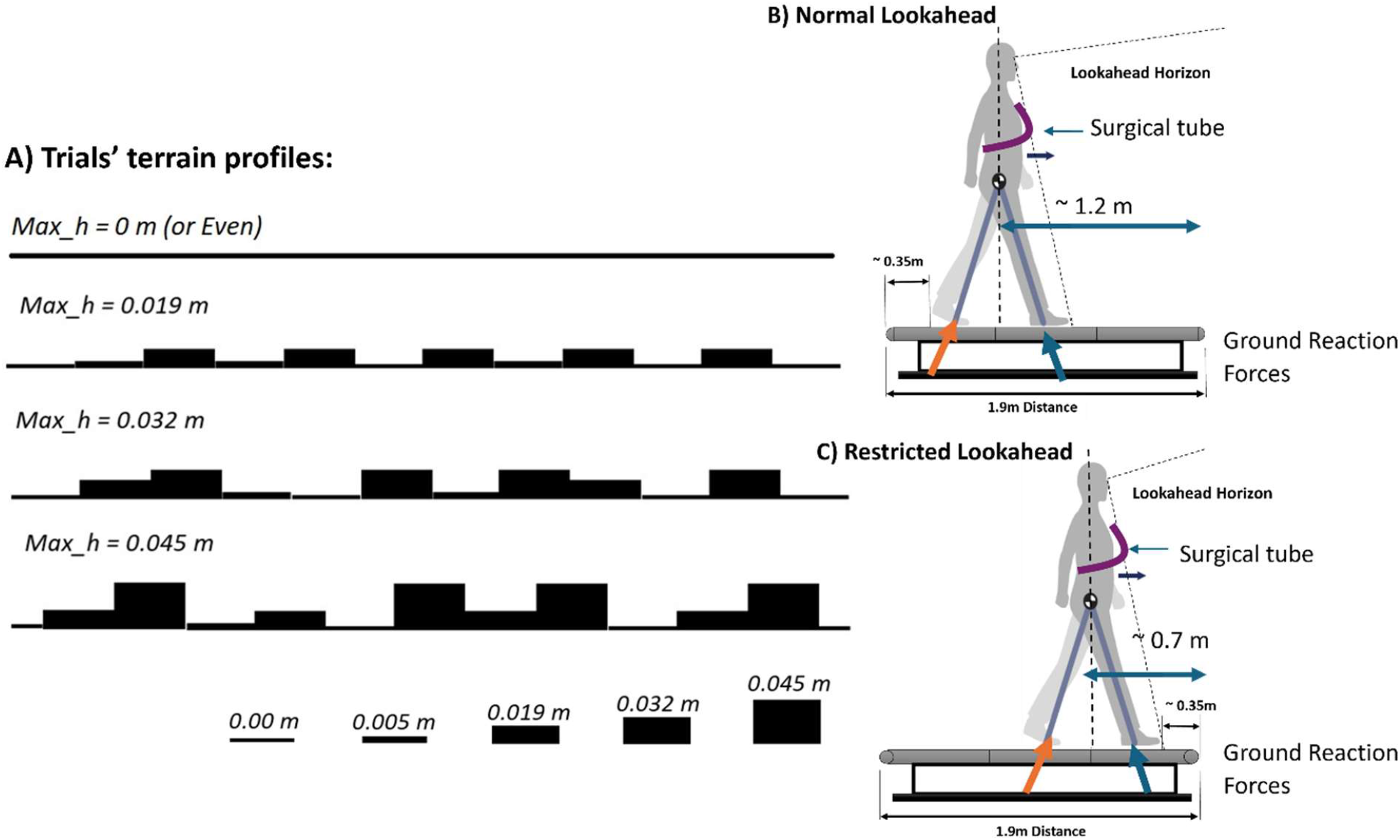
(A) The uneven terrain profiles were affixed across the surface of conventional treadmill belts. Each terrain was characterized by its maximum foam height (peak to peak: 0.019 m, 0.032 m, and 0.045 m). (B) Position for the normal lookahead. (C) Position for the restricted lookahead.

We placed subjects at 1.2m and 0.7m from the treadmill front to control their lookahead horizon. We adjusted each subject’s position during the initial trials at 1.2 m ⋅ s^−1^ to familiarize them with the experimental process. We called the first position Normal Lookahead, where subjects were asked to look at the coming terrain and walk normally (Figure *1*B). Likewise, we called the second position Restricted Lookahead (Figure *1*C). We asked subjects to walk naturally while they were looking forward. To assist subjects in maintaining their set positions, we mounted a surgical tube with some slack across the treadmill on two tripods. We asked subjects to touch the tube with their torso gently during the trials. We also provide verbal feedback when needed.

### Subjects

To include the effect of aging, we invited two groups of healthy young adults (age: 27.69 ± 5.21, mass: 71.0 ± 6.16 kg, leg length: ± 0.033m, six females and four males) and older adults (age: 66.1 ± 1.44, mass: 77.34 ± 12.13 kg, leg length: 0.935 ± 0.057, six males and four females). The subjects provided written informed consent before the start of the trials, and the University of Calgary Board of Ethics approved the experimental procedure.

### Experimental Result Analysis

When the walking speed was stabilized, we collected the Ground Reaction Forces (GRF) for 60 seconds at a 960Hz sampling rate. The The step-to-step regulation of gait parameters of interest for older adults with restricted lookahead: (A) Push-off regulation is provided for reference to indicate the duration of active work regulation. (B) Mid-flight active work (rebound) inducement. (C) Effective leg length control. (D) Step length modulation. The dashed lines represent the regulation for young adults during uneven walking with restricted lookahead for reference.

Presents the trial velocities and terrain amplitudes schedule. The order of velocities, terrains, and state of lookahead were randomized. We calculated the COM velocity and instantaneous COM power using the method outlined by Donelan et al. (high pass filter cut-off frequency = 0.256Hz to remove the drift)[15] and Kuo et al. [16].

**Table 1:** The trial schedule represents the combination of terrain amplitudes and walking speeds used for the experimental trials for young and older adults. Trials with normal lookahead are identified with uppercase letters, while trials with restricted lookahead are represented by lowercase letters.

| Terrains Amplitudes (m) | 0 | 0.019 | 0.032 | 0.045 |
| --- | --- | --- | --- | --- |
| Trials Velocities (m/s) |  |  |  |  |
| 0.8 |  |  | oa-ya/ YA-OA |  |
| 1 |  |  | oa-ya/ YA-OA |  |
| 1.2 | oa-ya/ YA-OA | oa-ya/ YA-OA | oa-ya/ YA-OA | oa-ya/ YA-OA |
| 1.4 |  |  | oa-ya/ YA-OA |  |

We also employed the marker arrangement used by Voloshina et al. [1] with minor modifications (active markers, Phase Space, CA, USA, 960Hz). The duration of motion capture was the same as the GRF data recording. We calculated the step elevation changes as the difference between the consecutive averages of the minimum vertical positions of markers defining the first and fifth metatarsals. We also defined the effective leg length as the distance between the marker identifying the greater trochanter and talus for each leg.

We assumed the rebound work [16] was the positive mechanical energy inducement during the mid-flight phase (Figure 2A) which happened after the step-to-step transition. We performed a regression analysis to investigate the impact of step elevation changes when subjects encountered a certain perturbation (terrain amplitude change). We included the elevation changes from four steps before to four steps after that encounter as independent variables. At the same time, the rebound works were the dependent variable:

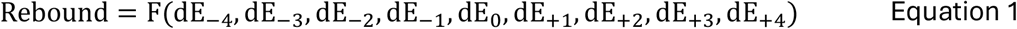

**Figure 2:**
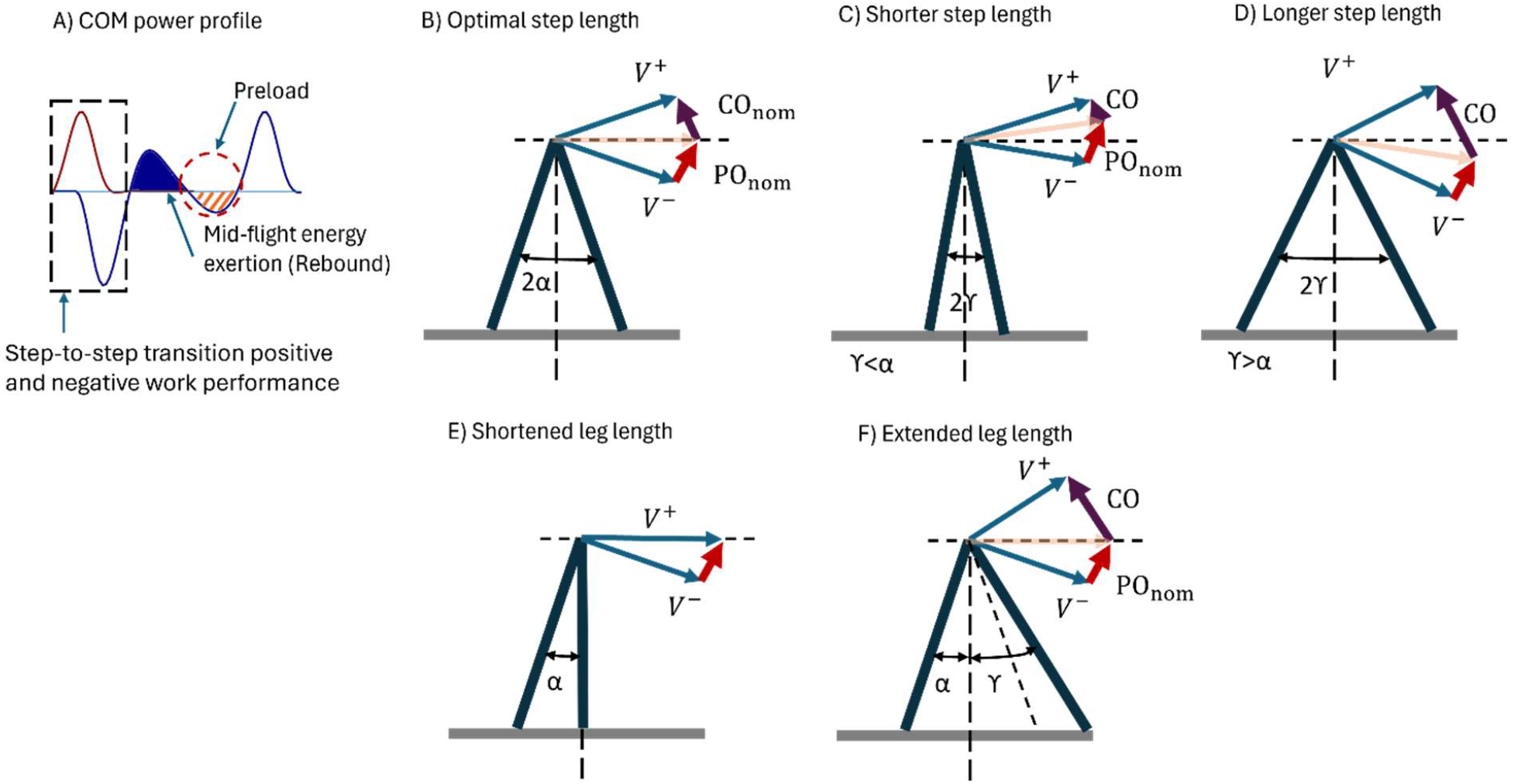
(A) The instantaneous COM power trajectory: the dashed box indicates the moment of the step-to-step transition, and the solid region (time integration) represents the mid-flight positive mechanical energy induction (rebound). The preload is suggested to absorb the rebound work. (B) Nominal step-to-step transition. (C) Step-to-step transition with nominal push-off and reduced step length. (D) Step-to-step transition with nominal push-off and longer step length. (E) Leading leg length shortening to accommodate a step up in which it landed vertically. (F) Leading leg extension; it was assumed that the upper bound change for leg extension was equal to the lower bound change for leg shortening departing from the nominal leg length.

In this case, the dE_i_ represented step “i” elevation change concerning a certain perturbation encounter. We considered the encounter to have happened at step “0.” We employed a similar approach for step length and effective leg length. Zero gains presented nominal values, while positive and negative gains were for values above and below the nominal magnitudes, respectively. We used Statsmodels 0.15.0 linear mixed effect model for statistical analysis. The significant results were reported as active modulation or control during the uneven walking.

### Simulation

We performed a simulation to quantify the effect of step length and effective leg variations on the step-to-step transition. First, for the average walking velocity of v = 1.25 m ⋅ s^−1^ that coincided with an average step length of 2α = 0.8 radiant [17], we altered the step length representing the angle between leading and trailing legs (2α) by ±15%. We assumed that the nominal push-offs were exerted [14]. Accordingly, we assessed the influence of the step length changes on the collision dissipation and the post-transition velocity (Figure 2B to D) [14].

Similarly, we assumed steps to start with nominal values for v = 1.25 m ⋅ s^−1^. Subsequently, we altered the leading leg length and maintained the COM elevation. We assumed that at the lower bound, the leading leg landed vertically. For the upper bound, we assumed the leg length would increase by the same amount as the vertical leg landing, which decreased the step length from its optimal velocity magnitude (Figure 2E & F) [18].

## Results

### Simulation Results

In the first simulation, for a walking speed of 1.25 m ⋅ s^−1^, the step length varied from 0.68 m to 0.98 m, and the nominal push-off impulse was 0.53 m ⋅ s^−1^. Accordingly, its associated active work was 0.14 J ⋅ kg^−1^. As the step length varied, the collision impulse and dissipation rose from 0.38 m ⋅ s^−1^ to 0.67 m ⋅ s^−1^ and from 0.07 J ⋅ kg^−1^ to 0.23 J ⋅ kg^−1^, respectively. On the other hand, the post-transition velocity ranged from 1.30 m ⋅ s^−1^ to 1.18 m ⋅ s^−1^ (Figure 3).

**Figure 3:**
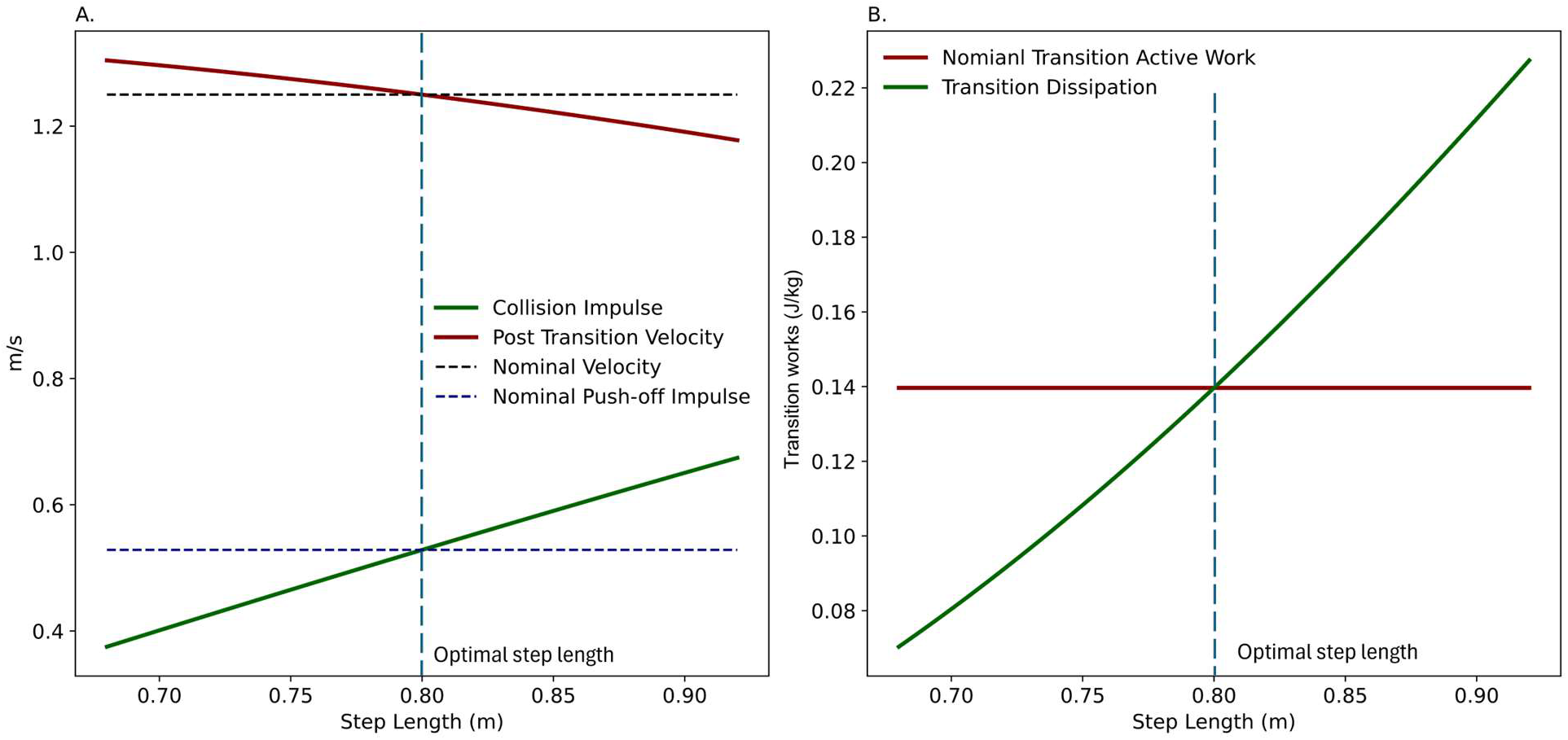
Simulation results for the nominal push-off work with step length variations: (A) Collision impulse and post-transition velocity varied as the step length changed. The dashed lines represent the nominal push-off impulse and velocity. (B) The nominal push-off and the collision dissipation changed as the step length was altered. It was assumed that the push-off was constant and nominal.

In the second simulation, the effective leg length varied from 0.92 m to 1.1 m (assuming the intact leg length = 1 m). Assuming the trailing leg maintained a nominal motion at the transition point (angle with vertical α=0.4 radian), the step length increased from 0.4 m to 0.98 m (small angle approximation [14], Figure 4). Subsequently, the collision impulse and its mechanical energy waste ranged from 0 m ⋅ s^−1^ to 0.74 m ⋅ s^−1^ and from 0 J ⋅ kg^−1^ to 0.26 J ⋅ kg^−1^. The post-transition velocity also declined from 1.36 m ⋅ s^−1^ to 1.14 m ⋅ s^−1^.

**Figure 4:**
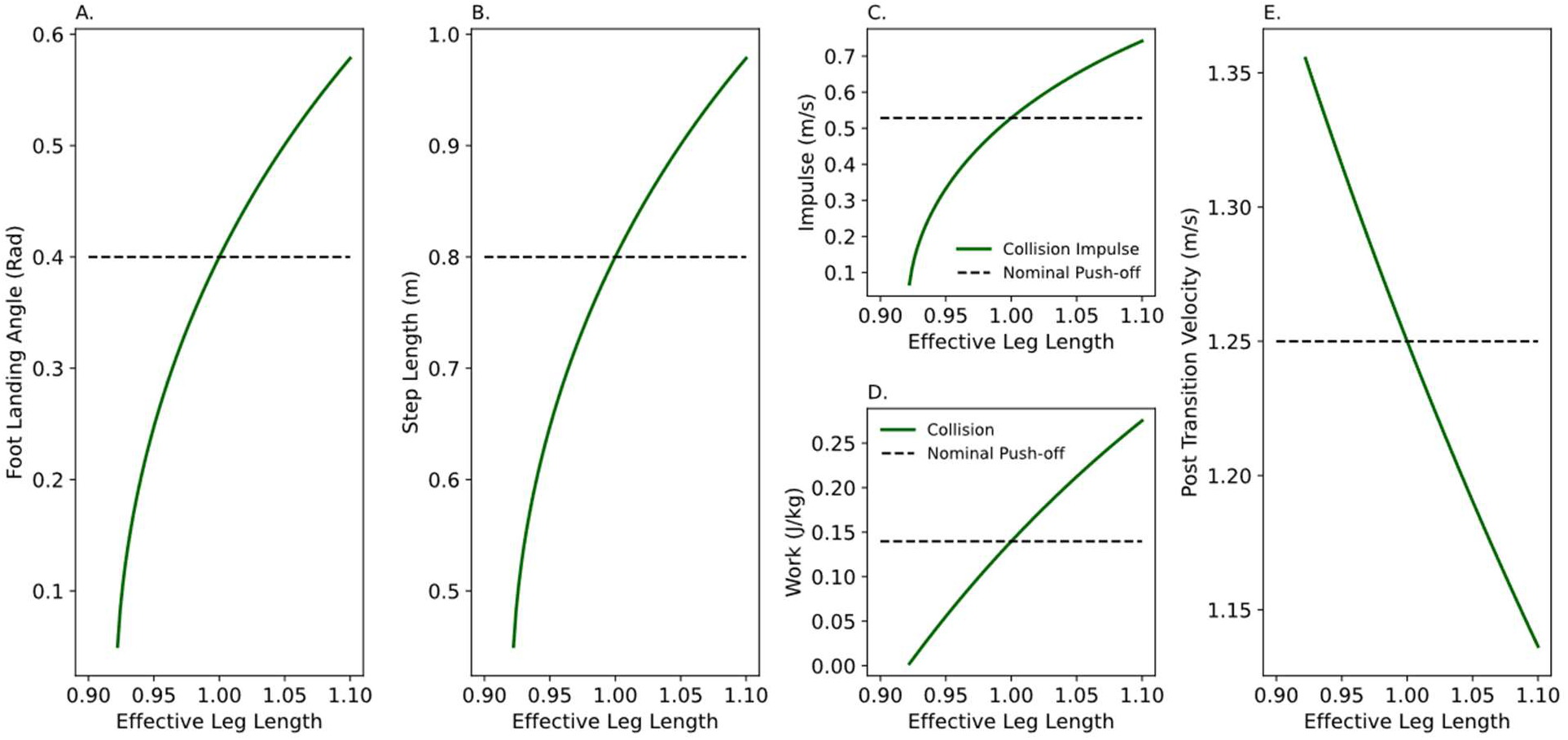
The effective leg length modulation and its consequences on the step-to-step transition dissipation. The COM elevation was assumed to be maintained at the nominal magnitude with the leg length change. (A) Leading leg foot landing angle with vertical. (B) Step length change with effective leg length variations. (C) Collision impulse variation with the effective leg length change after nominal push-off work induction. (D) Collision work dissipation change with the effective leg length change after the nominal push-off work exertions. (E) Post-transition velocity with effective leg length change.

### Experiments Results

The uneven walking affected the gaits parameters of interest (Figure 5). Younger adults with normal lookahead rebound works were lower than nominal at step -3 (gain = -0.81 J ⋅ kg^−1^ ⋅ m^−1^) and step -1 (gain = -0.85 J ⋅ kg^−1^ ⋅ m^−1^). At the point of encounter, the rebound rose substantially (gain = 1.45 J ⋅ kg^−1^ ⋅ m^−1^), while it dropped drastically when subjects left the perturbation (gain = -1.18 J ⋅ kg^−1^ ⋅ m^−1^). The step length was reduced at step -1 (gain = -0.31 m ⋅ m^−1^). The step length at the encounter and the subsequent step were raised and declined (gain = 0.35 m ⋅ m^−1^ and gain =0.29 m ⋅ m^−1^, respectively). The effective leg length periodically increased and declined with the terrains step downs and step ups (gains = -0.091, 0.097, -0.082, 0.096, -0.094, 0.082, -0.084, 0.081, and -0.083 m ⋅ m^−1^).

**Figure 5:**
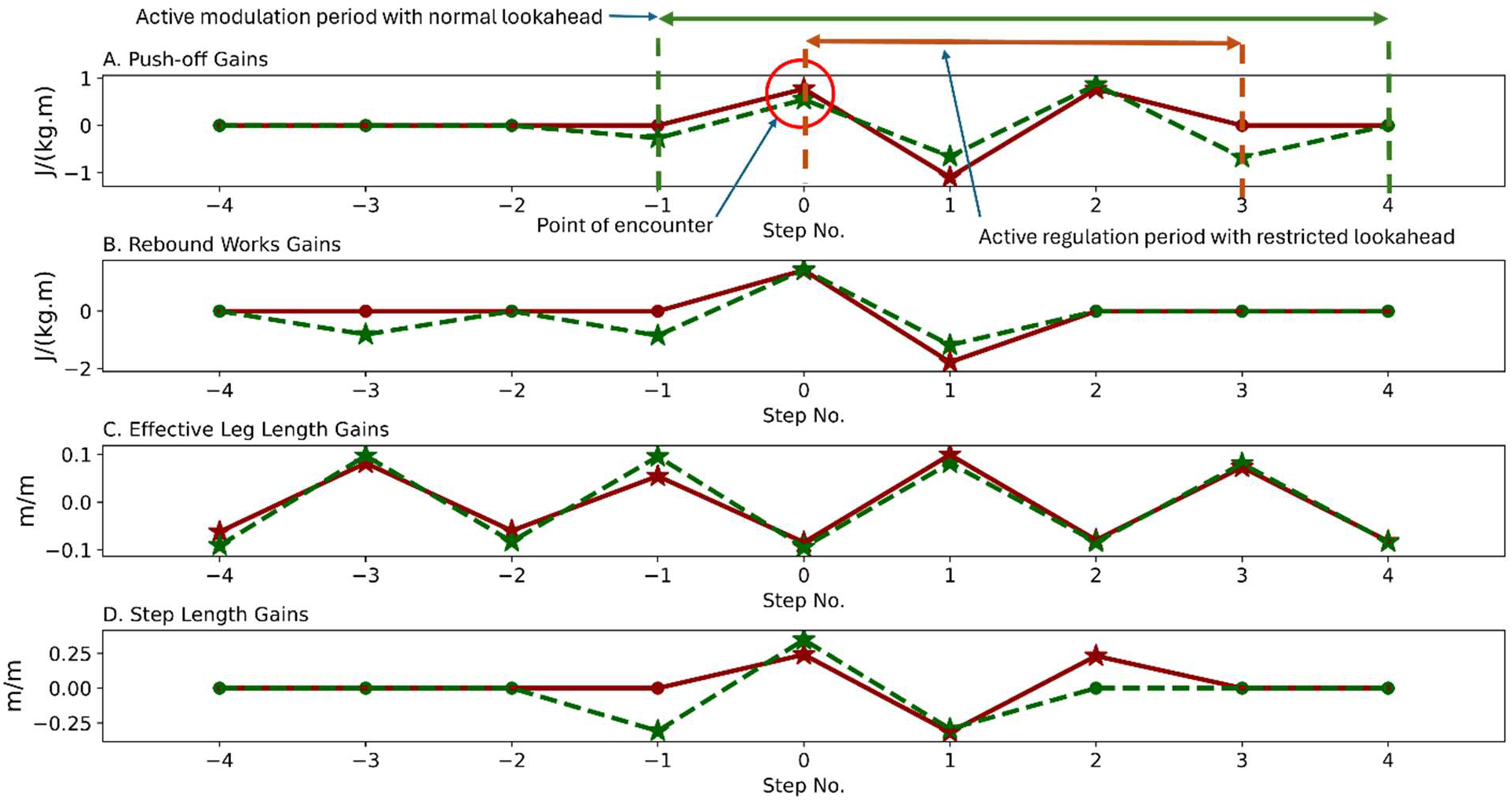
The step-to-step modulation of gait parameters of interest during uneven walking for young adults: (A) Push-off modulations are provided for reference to indicate the duration of active work modulation or regulation. (B) Mid-flight work (rebound) induction. (C) Effective leg length regulation. (D) Step length modulation. The dashed line shows the results for the normal lookahead traces, while the solid lines exhibit the restricted lookahead modulations.

Young adults with restricted lookahead mid-flight active modulations (rebound) were limited to the point of encounter and step +1, at which rebound work grew and declined drastically (gain = 1.43 J ⋅ kg^−1^ ⋅ m^−1^ and gain = -1.78 J ⋅ kg^−1^ ⋅ m^−1^, respectively). The step length stayed the same before the encounter. At the point of encounter, the step length rose (gain = 0.24 m.m^-1^), followed by a decline (step + 1 gain = -0.32 m ⋅ m^−1^) and an increase at step +2 (gain = 0.23 m ⋅ m^−1^). Like walking with the normal lookahead, the effective leg length increased and reduced periodically with step down and step-ups (gains = -0.062, 0.082, -0.060, 0.055, -0.084, 0.099, -0.079, 0.073, and -0.82 m ⋅ m^−1^) following the trial terrains profiles (Figure 5)

The older adults’ uneven walking differed in some key features from the young adults’ (Figure 6). With the normal lookahead, older adults demonstrated larger mid-flight and step length modulations, although the effective leg length modulation was similar. The rebound modulation exhibited an almost periodic pattern in all steps (fluctuations). The largest variations occurred at the point of encounter and step +1 (steps’ gains = 1.78, -1.62, 1.57, -1.53, 2.68, -2.81, 1.04, 0, and 2.22 J ⋅ kg^−1^ ⋅ m^−1^). The step length also fluctuated in all the investigated steps. Its oscillating trajectory gains were 0, -0.34, 0.44, -0.34, 0.39, -0.44, 0.32, 0, and 0 m ⋅ m^−1^). The effective leg length modulation seemed to match the trials terrains subsequent elevation changes (step-downs and step-ups, gains = -0.089, 0.084, -0.075, 0.086, -0.075, 0.075, -0.082, 0.076, and -0.089 m ⋅ m^−1^).

**Figure 6:**
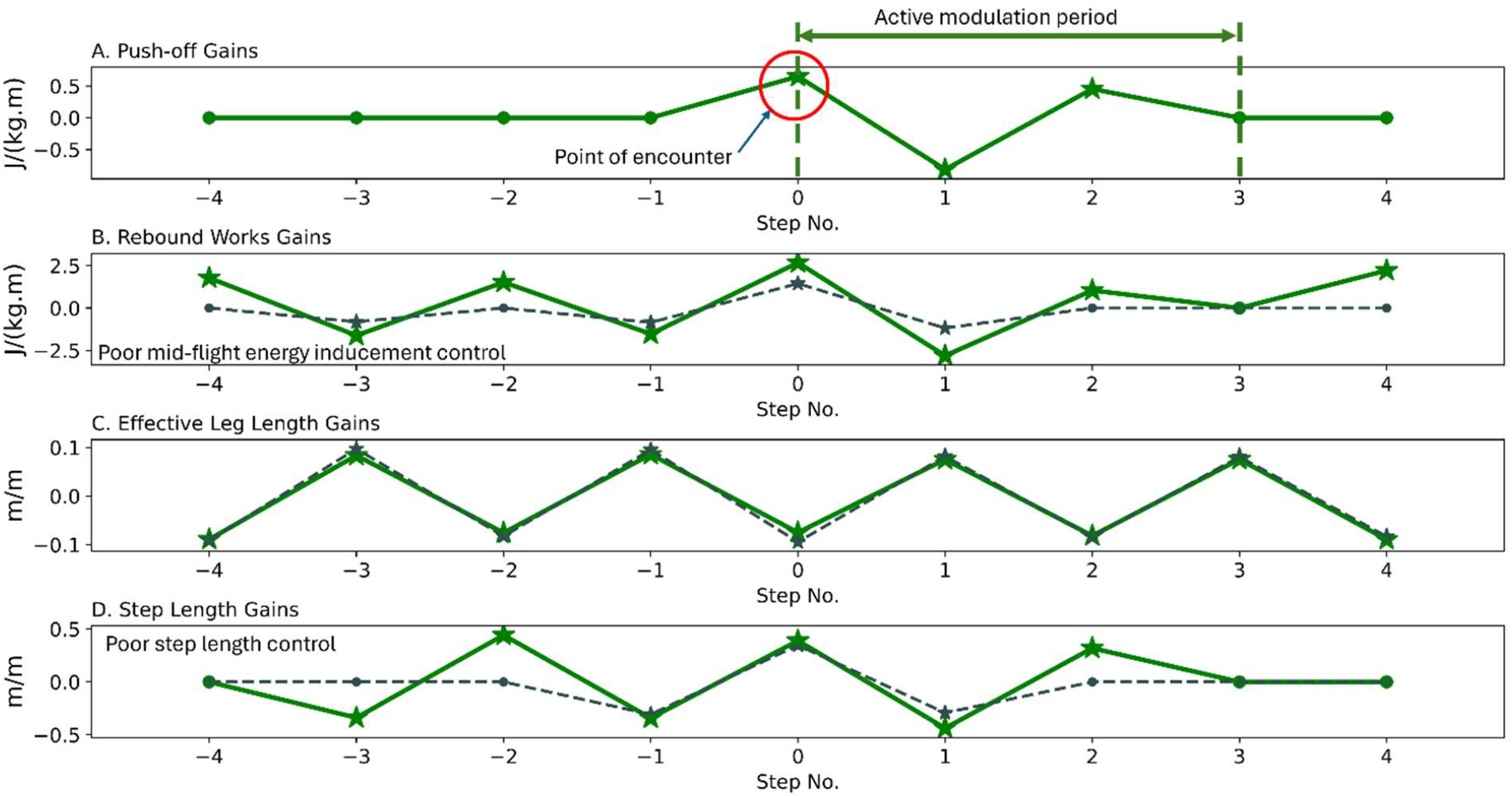
The step-to-step modulations of gait parameters of interest for older adults with normal lookahead: (A) Push-off modulation is provided for reference to indicate the duration of active work modulation. (B) Mid-flight work (rebound) inducements. (C) Effective leg length modulation. (D) Step length modulations. The dashed lines provide the modulations for young adults with normal lookahead for reference.

While older adults with the restricted lookahead, the mid-flight active work inducements (rebounds) were similar to those for young adults, their step length modulation was comparable to their trials with the normal lookahead (Figure 7). The mid-flight mechanical energy exertions were confined to the point of encounter and step +1 (gain = 0.93 J ⋅ kg^−1^ ⋅ m^−1^, and gain = -0.91 J ⋅ kg^−1^ ⋅ m^−1^, respectively). The step length regulated depicted oscillations before and after the point of the encounter (gains = 0, -0.28, -0.48, 0.66, -0.6, 0.32, 0, and 0 m ⋅ m^−1^). The effective leg length also depicted the oscillating pattern that went with the step downs and step ups of the trials’ terrains (gains = -0.12, 0.13, -0.14, 0.15, -0.16, 0.14, -0.14, 0.14, and -0.14 m ⋅ m^−1^).

**Figure 7:**
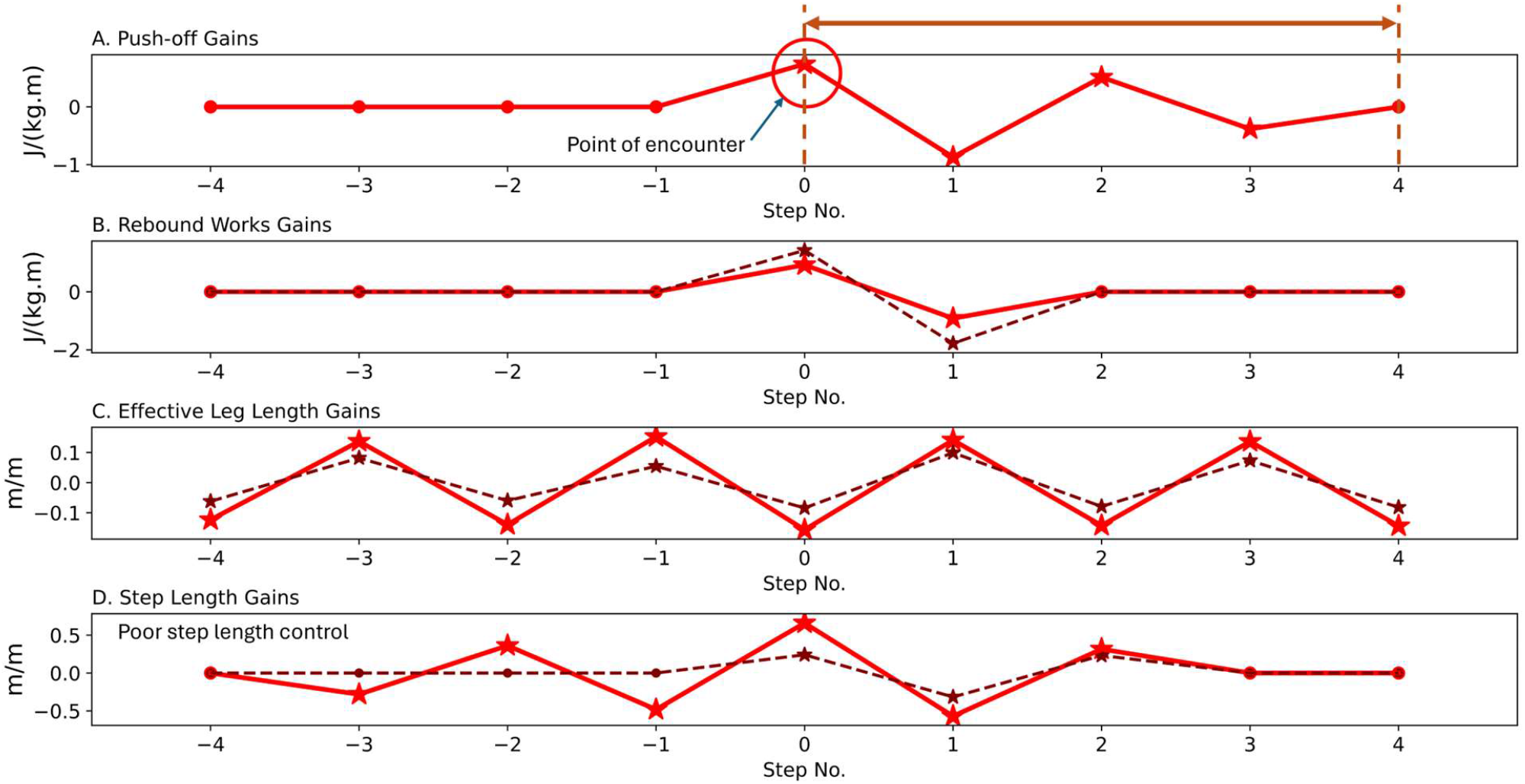
The step-to-step regulation of gait parameters of interest for older adults with restricted lookahead: (A) Push-off regulation is provided for reference to indicate the duration of active work regulation. (B) Mid-flight active work (rebound) inducement. (C) Effective leg length control. (D) Step length modulation. The dashed lines represent the regulation for young adults during uneven walking with restricted lookahead for reference.

## Discussion

Walking on uneven terrain presents challenges, notably an increase in energetic expenditure [19], [20]. Terrain complexities are quantified through terrain coefficients derived empirically [20], typically based on aggregate data from longer walking sessions [1]. Consequently, limited experimental work has focused on understanding step-to-step modulations [2], [5]. However, studying such modulations could offer insights into the control strategies humans employ before and after encountering terrain perturbations. This might reveal indications of efforts to limit mechanical work performance or conserve momentum. While previous studies have addressed active work and velocity controls through simulation [4], [5], [10] or experimental methods [5], our study aims to evaluate step-to-step modulation of effective leg length, step length, and mid-flight exerted mechanical works (rebounds).

Rebound work refers to the positive mechanical energy induction following the step-to-step transition [15], [16], [21]. It is theorized that rebound serves to straighten the stance leg and propel limb swings [16]. Additionally, it is suggested that rebound energy is nearly entirely absorbed by the preload during normal walking [15], [16] (Figure 2A). If the pre-emptive push-off [14] is insufficient to power the subsequent gait, additional active mechanical energy must be generated during the mid-flight phase [16]. Consequently, the magnitude of rebound must exceed the subsequent preload [16]. Conversely, if the push-off exceeds its associated collision dissipation or if the COM kinetic energy is elevated, such as through temporary COM elevation reductions [22] derived from potential energy, the subsequent rebound must diminish [16]. The decrease in rebound work observed prior to encountering perturbations in young adults with normal lookahead may result from excess COM kinetic energy that aligns with the prior perturbation dismounting. Conversely, a spike in rebound work atop the perturbation suggests a significant infusion of energy is required to partially counteract momentum loss. As subjects dismount from the perturbation at step +1, COM kinetic energy increases, thereby necessitating smaller active muscle work. The variance in rebound works between normal and restricted lookahead conditions may also stem from differences in push-off and other COM modulation controls.

The vertical leading leg landing (Figure 2E) established the lower bound for our effective leg length simulation. We utilized the magnitude of the resulting step length change from the nominal value (lower bound limit) to define our simulation’s upper bound, ensuring symmetry around the nominal magnitude (Figure 2F). Similarly, the experimental data also indicated comparable lower and upper bounds for leg length modulation (Figure 5C). This observation may suggest that humans tend to land their leading leg vertically during step downs [23], while extending the leg length within the confines of foot length and ankle range of motion [2].

Effective leg length may serve as a mechanism to adapt to perturbations in each step [2], [3]. Given that subjects can feel elevation changes in each step, ankle-sensitive and high-gain muscles [24] likely provide sensory information regarding the magnitude of step elevation changes. Our analysis indicates that legs are shortened in up steps and lengthened in down steps probably to minimize fluctuations in COM elevation. Consequently, leg length modulation may serve two distinct purposes during uneven walking. Firstly, at the point of encounter, reducing leg length limits the associated COM work against gravity [6]. When walkers dismount the perturbation, the reduction in COM elevation may result in excess kinetic energy that must be dissipated through increased collision at heel strike, potentially inducing high stresses on the limbs [3]. Therefore, after perturbation dismounting, extending the leg helps avoid excessive conversion of potential energy to kinetic energy.

However, at one step following the perturbation, landing the leading leg close to vertical, which shortens step length [23], limits collision dissipation and increases velocity post-transition to compensate for lost momentum. Yet, this strategy may lead to elevated metabolic costs [25]. Regardless of lookahead state, which may be a limiting factor in accommodating perturbations, effective leg length trajectory remains consistent across terrain amplitudes. Similarly, larger leg length modulation in older adults with restricted lookahead may further elevate energetic costs during uneven walking compared to young adults.

The simulation reveals that, for a given nominal positive active work (pre-emptive push-off), the COM velocity increases or decreases depending on whether the step length is shortened or lengthened. It is demonstrated that, for a constant speed, increasing step length amplifies transition mechanical work and its associated metabolic cost by the fourth power of the step length [15]. Consequently, increasing step length could be employed as a transient strategy to dissipate mechanical energy and slow walking speed. Conversely, reducing step length enhances walking velocity by reducing the transition energy dissipation [14], [15], [21]. Consequently, walkers may augment their walking momentum at the expense of metabolic energy, which may be lower than the equivalent mechanical work energetics. However, consistently extending shorter step lengths throughout walking elevates walking energetics by the fourth power of step frequency [26], [27]. It is proposed that at higher limb motion frequencies, the energetic cost of walking is primarily influenced by the force generation rate [18], [26], setting a lower limit for step length reduction.

During uneven walking, transient variations in step length may be categorized as active or passive. Studies indicate that during step-downs, step length is shortened, with the leg landing closer to vertical [23], while in step-ups, it tends to be longer [2], [3]. Consequently, with normal lookahead, the shorter step length detected before the encounter likely increases COM velocity just prior to the perturbation, enhancing the walker’s momentum (Figure 5D). Thus, such step length changes may represent active modulation. Conversely, both with normal and restricted lookahead, step length variations at and following the perturbation appear to be passive outcomes of step transitions. With restricted lookahead, the increase in step length at step +2 enhances collision dissipation, potentially representing another active regulation to reduce walking speed by dissipating energy.

In old-age walking with normal lookahead, fluctuations in active control are shown (Figure 6B & D). These fluctuations may be linked to the state of the Central Nervous System (CNS), which tends to deteriorate with age [28]. Consequently, the observed oscillations and overshoots may indicate diminished feedforward control in older adults. Conversely, with restricted lookahead, active work exertions do not exhibit noisy controls, but older adults’ step regulations do. The variability in step length modulations may also be influenced by the processing of other sensory information, such as touch, that older adults receive at each step contact (Figure 7D). Hence, it is plausible to infer that older adults’ feedforward control may be inferior to their feedback control.

We have observed that different gait parameters exhibit varying modulation time durations, with the condition of trials impacting them. For instance, with restricted lookahead, young adults’ mid-flight (rebound) modulation is limited to two steps, whereas step length is modulated over three steps (Figure 5B & D). Besides the order or amplitude of modulations, the lookahead state may also influence regulation duration when control transitions from feedforward (anticipatory) to feedback. For example, with normal lookahead, mid-flight modulations occur over four steps, whereas with restricted lookahead, the regulation period spans two steps (Figure 5B). Age likely has a significant impact, as control performance is influenced by processing and sensor states. Diminished performance of the CNS with age may similarly affect human sensory capabilities, potentially weakening the quality of sensory estimates. Moreover, the terrain complexity, which affects the frequency and amplitude of perturbations, may also impact input signals and CNS processing. Additionally, since control is executed through muscles and joints, their state [29], [30] may also influence the performance of older adults.

In summary, our observations reveal that in addition to the pre-emptive push-offs exerted at the start of each step, humans also exert muscle works during the mid-flight phase. Furthermore, various gait aspects may be altered actively or passively to regulate walking momentum. Step length exhibits both active and passive modulation changes, whereas effective leg length tends to remain consistent and actively regulated. The state of lookahead also influences regulation, whether through feedforward or feedback controls. We can identify indicators of anticipatory (feedforward) and feedback controls that change depending on the state of lookahead. It appears that the influence of lookahead state is significantly greater than other sensory information. Younger and older adults seem to respond differently to terrain complexities. Younger adults primarily concentrate active controls around the point of encounter, while older adults demonstrate poorer and more prolonged active control durations. Additionally, it appears that older adults have inferior feedforward control performance compared to feedback control.

## Acknowledgements

This work was supported in part by the Natural Sciences and Engineering Research Council of Canada (NSERC) Discovery and Canada Research Chair (Tier 1) programs.

## Notes

**Conflict of Interest:** None

### Competing Interest Statement

The authors have declared no competing interest.

